# Early fate of Exogenous promoters in *E. coli*

**DOI:** 10.1101/710772

**Authors:** Malikmohamed Yousuf, Ilaria Iuliani, Reshma T. Veetil, Aswin Sai Narain Seshasayee, Bianca Sclavi, Marco Cosentino Lagomarsino

**Affiliations:** LBPA, UMR 8113, CNRS, ENS Paris-Saclay, 61 Avenue du President Wilson, 94235, Cachan, France; National Centre for Biological Sciences, Tata Institute of Fundamental Research, Bangalore 560065, Karnataka, India; Sorbonne Université, Campus Pierre and Marie Curie, 4 Place Jussieu 75005, Paris, France; CNRS, UMR7238, 4 Place Jussieu 75005, Paris, France; IFOM, FIRC Institute of Molecular Oncology, Via Adamello 16, 20143, Milan, Italy; Physics Department and INFN, University of Milan, Via Celoria 16, 20133, Milan, Italy

## Abstract

Gene gain by horizontal gene transfer is a major pathway of genome innovation in bacteria. The current view posits that acquired genes initially need to be silenced and that a bacterial chromatin protein, H-NS, plays a role in this silencing. However, we lack direct observation of the early fate of a horizontally transferred gene to prove this theory. We combine sequencing, flow cytometry and sorting, and microscopy to monitor gene expression and its variability after large-scale random insertions of a reporter gene in a population. We find that inserted promoters have a wide range of gene-expression variability related to their location. We find that high-expression clones carry insertions that are not correlated with H-NS binding. Conversely, binding of H-NS correlates with silencing. Finally, while most promoters show a common level of extrinsic noise, some insertions show higher noise levels. Analysis of these high-noise clones supports a scenario of switching due to transcriptional interference from divergent ribosomal promoters. Altogether, our findings point to evolutionary pathways where newly-acquired genes are not necessarily silenced, but may immediately explore a wide range of expression levels to probe the optimal ones.

## 1 Introduction

The high fraction of mobile genes in bacterial genomes is a source of a great diversity of phenotypes. This large diversity challenges the very concept of species, and has enormous importance for understanding pathogenicity and antibiotic resistance [1]. At the genetic level, *E. coli* genomes vary dramatically in their sizes ranging from 4.5Mb to 6Mb. Comparative genomic surveys of *E. coli* have shown that there is a core set of genes which is highly conserved across the species and coexists with a large pangenome, the set of genes that can be gained by horizontal gene acquisition from other species [2]. Indeed, bacteria acquire exogenous DNA by transformation (of naked DNA from the environment), transduction (of DNA from bacteriophages) or conjugation (from fellow bacteria through molecular pipes such as pili) [3]. In order to be functional, exogenous acquired genes often need for the metabolic and the regulatory circuitry of the cell to be rewired [4, 5]. Furthermore, expression of a foreign gene can interfere with the resources allocated for endogenous gene expression. Therefore, horizontally acquired genes must be regulated [6].

A primary mode by which the expression of horizontally-acquired genes is regulated is believed to be transcriptional repression, which is achieved by proteins such as H-NS in enterobacteria, including *E. coli* [7, 8, 9, 10](reviewed in ref. [11]). Many previous studies support both the need of initial repression of acquired genes, and the view that H-NS repression is relevant for the successful establishment of these genes [12, 5, 13, 6]. H-NS is among several “global” transcriptional regulators that affect the expression of hundreds of genes in *E. coli*. It binds to AT-rich or intrinsically bent DNA sequences and forms structures such as stiff rods or DNA-protein-DNA bridges, which might act as geometrical motifs for transcriptional silencing [14, 15]. Many H-NS binding regions are up to a few kilobases long. The length of these binding regions correlates with the degree of transcriptional repression imposed on the target gene [9, 16] and genes regulated by H-NS are very highly expressed in the absence of this repressive control [17, 18]. Since the levels and activity of H-NS depend on environmental conditions and growth rate, this level of regulation allows for a coordinated gene-expression change needed for cellular adaptation. At least through the action of H-NS, which is a notorious nucleoid-shaping protein [19], the dynamics of horizontal transfers is related to the physical organization of the chromosome. An important question to be addressed is whether and how the organizational features of the *E. coli* chromosome (such as the “macrodomain” architecture [20, 21, 22, 23] are correlated with gene acquisition and control of gene expression, particularly of acquired genes [24].

Horizontally transferred genes are often clustered along the genome [25, 26, 22, 27, 3]. In part, this reflects joint transfer of functionally co-dependent genes that would provide no benefit if transferred independently. In part, however, this reflects the existence of “permissive” zones along the chromosome, which experience recurrent integration and high turnover. Permissive zones can originate or be reinforced through physical integration biases, where the presence of integrases and/or recombinogenic sites facilitates acquisition of genetic material [1]. Additionally, in many species including *E. coli*, horizontally acquired genes preferentially accumulate near the (AT-rich) terminus region [2, 22, 1], possibly to avoid deleteriously high expression near the origin due to gene copy number effects.

The genome sequence organization of a given species is a result of selection pressure and architectural constraints. Some of these have been clearly identified [17, 3, 28, 27, 6]. For example, highly expressed, newly acquired genes must be kept from interfering with the expression of essential genes. However, comparatively little is known about the early dynamics of acquired genes. Do clear physical insertion biases emerge? What are the phenotypic impacts of inserted genes and how are they linked with expression levels? How, in turn, is gene expression a consequence of the locus of insertion? Are they immediately silenced and do H-NS and nucleoid organization play a role?

To access some of the above questions, we devised an experimental assay (Fig. 1) where a cassette including an antibiotic resistance gene and a GFP reporter under control of a highly expressed ribosomal promoter is inserted systematically in the genome, and the resulting mixed and clonal populations are analyzed by sequencing and single-cell biology methods. This methodology allows us to describe statistical tendencies for a reference promoter to be inserted and initially maintained in specific chromosomal contexts, as well as to characterize its fate in terms of both gene expression activity and noise of the transcription reporter constructs at different insertion sites on the genome.

**Figure 1:**
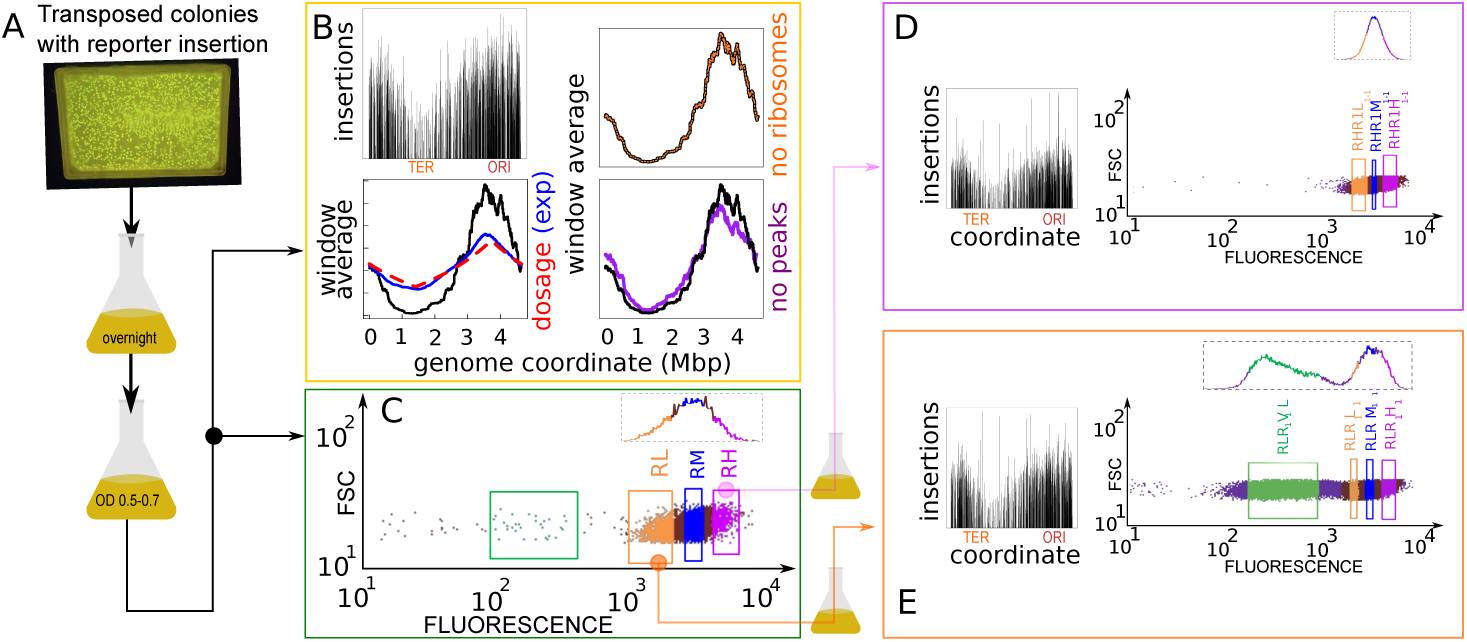
Insertion localization and sorting by gene expression. A: Experimental pipeline. Massive transposon insertion of a GFP reporter gene cassette in around 100000 founder strains was tested by plating on kanamycin-selective agar and PCR. Surviving colonies were mixed, grown overnight in LB, resuspended and grown to a fixed OD. B: Sequencing of resulting parental populations yields the locations of the insertions. The bottom panel compares a 3kb sliding average of the coverage (black line, y axis rescaled for comparison) with the prediction from gene dosage, and the experimental dosage (red dashed line) measured by whole-genome sequencing (blue line) the right panels are controls that the trend of insertions copy number is not due to ribosomal genes and (orange line) and to the insertions with top 10% coverage (> 3000 reads/bin, purple line). C,D,E: forward scatter vs GFP expression measured by flow-cytometry. FACS Sorting by the level of fluorescence was performed on a total of four rounds (see Supplementary Fig. 1). Selecting for high expression (RH) yielded a population with a similar distribution of gene expression (C), while selecting for low expression yielded a population with a bimodal distribution of gene expression (D). Insets show insertions found by population sequencing (y axis are counts in logarithmic scale), with overall similarity but local differences.

## 2 MATERIALS AND METHODS

### 2.1 Experimental procedures

#### Generation of the transposon library

The transposon used for genome wide insertion is derived from Ez-Tn5*™* custom transposon kits (Epicentre). The insert is a cassette composed of the resistance gene to kanamycin and the *gfpmut2* gene under control of the well characterized ribosomal promoter rrnBP1. The rrnBP1 promoter is sensitive to both changes in DNA supercoiling and to regulation by the small metabolite ppGpp. The full length rrnBP1 promoter, “P1-long”, includes binding sites for H-NS and Fis. We also used as a control a shorter version “P1-short”, which lacks the Fis sites and has reduced H-NS binding [29].

The level of expression of GFP under control of rrnBP1 can be thus used to assess the effect of nucleoid associated proteins such as H-NS.

The cassette is cloned between the BamH1 and Sal1 sites of the Ez-Tn5*™* pMOD*™* -3 construction vector. The 5’ phosphorylated primers used for amplifying the insert with mosaic ends of the transposon are the following,

5-CTGTCTCTTATACACATCTCAACCATCA-3 5-CTGTCTCTTATACACATCTCAACCCTGA-3

200 nanograms of the purified PCR amplified fragments were mixed with the transposase enzyme and glycerol at 37*°C* for 1 hour. Bacteria were grown to early log phase in LB at 37*°C* before the transposons were added. The transposon mixture (1*µ*l) was electroporated into the wild type strain (BW25113). After electrotransfromation, the bacterial cells were allowed to recover in SOC (rich) medium at 37*°C* for 2 hours without antibiotics. The cells were plated on kanamycin (50*µ*g/ml) plates, and grown overnight at 37 degrees to select the transformants. Genomic DNA was isolated from the recombinant clones and internal primers binding to the transposon-specific regions were used to verify the insertion by PCR. A typical electroporation experiment yielded 15,000 to 20,000 transformants. In total, approximately 10^5^ transformants were pooled from 10 identical electroporation experiments. These are called the parental clones or the parental population in the main text.

#### Sorting of transformed clones using FACS

The transformed cells were pooled together from the Kanamycin agar plate and were grown overnight at 37*°C* in minimal medium (0.5% Glucose + 0.2% CAA). On the following day, 1% of the primary culture was diluted to 1:500 and grown in a secondary culture in minimal medium until the OD reached 0.5 to 0.6 (mid exponential phase). Different populations were sorted by FACS (Fluorescent Activated Cell Sorting) by the level of GFP expression. The populations in the left and right tail of the distribution (roughly below and above 0.6 standard deviations from the mean) after the first round of FACS, and those around the mean of the distribution (roughly 0.5 standard deviations from the mean) were sorted and labelled (see Fig. 1 and Supplementary Fig. S1 for a graphical scheme) as low (RL), medium (RM) and high (RH) expression respectively (R stand for Rounds of FACS). Subsequently, the RL sorted population underwent a next round of FACS to yield 4 sorted subpopulations, RLR1V1L1 (very low), RLR1L1 (low), RLR1M1 (medium) and RLR1H1 (high) whilst the RH yielded three populations, RHR1L1-1 (low), RHR1M1-1 (medium), RHR1H1-1 (high). A third round of FACS was performed on RLR1V1L1 (very low), RLR1L1 (low), and RLR1H1 (high) selected from the low expressing population, and on two populations, RHR1L1-1 (low) and RHR1H1-1 (high) from the high expressing RH population. Only the very low expressing population, R1L1R2V2L2 from the third round of FACS underwent a fourth round of FACS resulting in four subpopulations that were sorted depending on their levels of expression. All the populations from the four rounds of FACS were stored at -80^*o*^ C in Glycerol.

#### Library preparation and sequencing of the different populations

##### Tradis sequencing for FACS sorted populations

Bacterial populations sorted based on GFP fluorescence from FACS experiments were grown in LB media containing the antibiotic Kanamycin for 16 hours at 37*°C* at 200 rpm. The genomic DNA samples were isolated from these populations using the QIAGEN DNA purification kit protocol. Genomic DNA was sonicated (Covaris S220) to 350 bp fragments using standard factory settings, was end-repaired using Truseq Nano DNA LT End Repair and A-Tailing mix. The end-repaired samples were ligated using Truseq Nano DNA LT ligase enzyme to adapters described by ref. [30]. Adapter-ligated fragments were purified and PCR was performed to enrich the fragments. The PCR primers were designed according to ref. [30]. For the PCR, we used Truseq Nano DNA LT PCR mix with thermal cycler conditions of 95*°C* for 3 min followed by 19 cycles of 98*°C* for 20 s, 65*°C* for 30 s, and 72*°C* for 30 s, with a final extension of 5 min at 72*°C*. PCR products were purified with 1 AMPure XP beads (Agencourt) and quantified by Qubit. The pooled library contained three samples at 6 pM with 9% PhiX and was sequenced with a 50-cycle single-end MiSeq reagent kit (Illumina). The sequencing reads were aligned to the reference genome (NC 000913.3) using the BWA (Burrows-Wheeler Aligner) method. Mapped reads were binned in to 10 kb non overlapping bins using a custom perl script and the number and frequency of insertions in each bin plotted against the chromosome coordinates. The reads considered were filtered for map quality greater than 20 from the same file.

##### Whole genome sequencing to measure gene dosage

For genomic DNA extraction, the overnight cultures were inoculated in 50 ml of fresh LB media to bring the initial OD of the culture to 0.03 and the flasks were incubated at 37*°C* with shaking at 200 rpm. Cells were harvested at the maximum growth rate and genomic DNA was isolated using SIGMA GenElute*™* Bacterial Genomic DNA Kit using the manufacturer’s protocol. Library preparation was carried out using the Truseq Nano DNA low throughput Library preparation kit and Paired end sequencing of genomic DNA was performed using Illumina Hiseq 2500 platform. The sequencing reads were aligned and mapped to the reference genome (NC 000913.3) using Burrows Wheeler Aligner (BWA) specifying alignment quality and mapping quality thresholds as 20. Read coverage across the genome was calculated for non-overlapping windows of 200 nt each using customized perl scripts.

##### Whole genome sequencing and analysis of single clones isolated from the FACS populations

Genomic DNA of single clones isolated from sorted populations (Qiagen DNA purification kit) were subjected for paired-end sequencing (2X75) using Illumina Miseq platform. DNA libraries were prepared using the TruSeq Nano DNA LT kit protocol. The sequencing reads were mapped with the reference genome (NC 000913.3) as well as with the known cloned insertion sequence using the BWA (Burrows-Wheeler Aligner) method. The positions of the insertions were identified in each clone using the mate pairing approach for paired-end reads in which the cloned insert sequence is mapped to the read where the genome coordinate maps to the pair.

##### Nanopore sequencing to confirm the transposon insertion in a rRNA operon

A total of 32 samples were shown to have the insertion in a rRNA operon from whole genome sequencing data. Five of these were subjected to nanopore sequencing using Oxford nanopore technologies. Reads were assembled using a *de novo* assembler [31] to obtain a single contig(∼4.6 Mb) for each sample. A BLASTN search was carried out using the cloned insert sequence as a query to confirm the chromosomal position of insertion.

#### Plate reader analysis

The growth media used in this study were a “slow growth” medium, M9 minimal medium + 0.5% glucose and “fast growth” medium, M9 minimal medium + 0.5% glucose + 0.2% casaminoacids. All the strains were grown overnight in M9 medium + 0.5% glucose + 0.2% casaminoacids. The next day, they were diluted to 1:1000 for the slow growth medium and 1:10000 for the fast growth medium in a flat bottom 96-well plate. The final volume in each of the culture well was 150 *µ*l. 70 *µ*l of mineral oil (Sigma) was added on top to reduce evaporation of the samples. The plates were incubated at 37*°C* shaking in the plate reader (Tecan). The optical density (OD610) and the GFP fluorescence at 535 nm were measured every 5 minutes over 21 hours. As a control strain, BW25113 was used, not containing the GFP reporter insertion in the chromosome.

#### Data analysis of the plate reader experiments

Growth rate and GFP concentration for all the strains were obtained from the data obtained in the plate reader on the optical density at 610 nm (OD) and the fluorescence (GFP) as a function of time. The exponential growth phase is defined as the period during which the OD increases exponentially between two user-defined OD thresholds. The growth rate in exponential phase is defined as the slope from the linear fit of the log(OD) vs time divided by log(2). The GFP concentration is obtained from the slope of the linear fit of the plot of GFP versus OD.

#### Measurement of the noise in gene expression

30-32 clones from each population (a total of 658, plus the controls) were selected randomly from the agar plates. After overnight primary growth in M9 minimal medium with 0.5% Glucose and 0.2% casaminoacids, 1% of the primary culture was diluted to 1:500 and grown in a secondary culture in the same medium until the OD reached 0.5 to 0.6 (mid-log Phase). When the cultures reached mid-log phase, we performed FACS analysis to measure the distribution of fluorescence. 50,000 events were collected for each clone. The first 1000 events were omitted from further analysis. Using the FlowJo software, a kernel density estimate was used to select the contour with the highest density of cells and the center region of the contour was taken for analysis. The magnetic gate plug-in from FlowJo was used to selectively encompass 10,000 events, and the gate was re-adjusted if there was a shift, so that only the most dense centered region events (10,000) were collected for further analysis. The same gate was used for all the clones in a 96-well experiment. A custom-made MATLAB code was used to measure the coefficient of variation (CV), Fano factor, and noise of each sample.

#### Time-lapse, single cell imaging of the clones and data analysis

Individual colonies were grown overnight in minimal medium containing 0.5% Glucose + 0.1% CAA. The overnight culture was diluted to 1:200 in a secondary culture in the same growth medium. The culture was grown until the OD reached 0.1.

##### Agar-pad Protocol

Five microliters of culture were added to the center of a 1.25% (w/v) low-melting agar pad made with the same growth medium as the culture. The pad was allowed to dry for half an hour at room temperature and then it was inverted into a small glass dish to be observed under the microscope.

##### Microscope

A Nikon Inverted microscope ECLIPSE Ti-E with a 100X oil objective was used to perform the live imaging. To correct the drift in focus, we used a Nikon Perfect Focus System. A custom-made temperature control system was used to keep the temperature constant at 30*°C* throughout the imaging session. The automated x-y stage was used to capture different regions of interest in the agar pad. Time-lapse movies were captured over different positions at intervals of 5 minutes.

##### Exposure time and Acquisition parameters

For the repressed or silent colonies, the settings were following: exposure time: 200ms, multiplier: 100, lamp 20%. For the low noise colonies, the settings were following: exposure time: 40ms, multiplier: 10, lamp 15%. The reason for the difference is that the signal from low-noise colonies saturated with the settings of high-noise ones, whose signal can be very low.

##### Data analysis

The contours of the cells were drawn manually in every frame using the ImageJ software. Total GFP intensity was measured for every cell in the lineage tree. The total intensity was divided by the area to obtain the mean fluorescence intensity of each cell in the frame. The background intensity was subtracted to obtain the normalised fluorescence intensity of the cells.

#### Data sets from the literature

The RegulonDB database (http://regulondb.ccg.unam.mx/) was used to extract gene positions. The lists of genes tested for insertion enrichment were obtained from the NuST database [32], which collects data from many published studies related to nucleoid organization[32]. The complete list of data sets is presented in Supplementary Table 1.

Each data set has the form of a gene list. The data sets were organized by the type of biological data and the experimental techniques, and divided into the following wide categories: “Genes linked to specific phenotypes”; “Genes next to binding sites of a regulator from binding data”; “Genes sensitive to nucleoid perturbations from transcriptomics experiments”; “Genes with specific annotations”; “Mutated genes in driven evolution experiments”; “Target genes of a regulator”. A description of the gene lists used in the analysis is provided in Table 1.

The OriC site is the origin of replication of the *E. coli* chromosome. For historical reasons, the coordinates of the *E. coli* circular chromosome do not start at the origin of replication, but at the origin of transfer during conjugation. We set the OriC position at [3,925,744 - 3,925,975] (https://ecocyc.org/)

## 3 RESULTS

### 3.1 Efficient protocol for production and characterization of systematic exogenous reporter insertions

The promoter chosen to control GFP expression in the randomly inserted cassette is the rrnBP1 promoter of the rrnB ribosomal operon. We chose this well-characterized highly expressed promoter because it is regulated by changes in DNA supercoiling and by the abundant nucleoid proteins Fis and H-NS [29]. We also considered, as a control, a shortened version of the promoter lacking regulation by the nucleoid proteins Fis and H-NS but still regulated by DNA topology.

Fig. 1A describes our pipeline (see methods and Supplementary Fig. S1). On the order of 10^5^ transposed colonies (this estimate was based on manual counting as we knew the number of transposed colonies in each plate) were mixed and grown overnight in minimal medium. This population was regrown to a fixed OD in the same medium, and initially assayed by population sequencing (Fig. 1B and flow cytometry Fig. 1C,D,E). We then used a cell sorter to select sub-populations based on gene expression levels (Fig. 1C,D,E). Finally, 658 randomly hand-picked clonal populations were individually characterized by flow cytometry. A subset of 96 from the 658 clonal populations were sequenced, and 90 of these were used to measure gene expression and growth rate in a plate-reader assay. A smaller selected subset of clones was used to measure the dynamics of gene expression in single-cell microcolony growth assays by epifluorescence microscopy.

### 3.2 Bimodal distribution of gene expression in parental populations and low-expressing sub-populations

Comparison of the fluorescence distribution in the sorted populations obtained from the high (RH) and low-expressing (RL) fractions of the parental population (Fig 1C,D,E) shows that the low-expressing (RL) population have a sub-population of clones with very low expression. In the parental population (Fig 1C), some of these low-expression clones are already visible (green box). Sorting them from the RL population gave rise to the population RLR1V1L.

Outside of this low-expression peak, the distribution of gene expression has little variation in the sorted populations compared to the parental one. This variability is the combination of the variability of promoter expression across single cells that are clonal (i.e. where the insertion is in the same exact position) and the variability of mean expression between clones with different insertion locations. Thus, the clonal variability should be considerably high in order to account for the fact that the overall pattern of variability is robust in the sorted sub-populations (which contain less clonal variants). There are, however, some important differences in the distributions, mirrored by differences in the location and frequency of the insertions in the sorted populations, which turn out to be significant (see below).

### 3.3 Insertions are non-uniform and sparse and are more biased towards the replication origin than justified by gene dosage

TraDIS Sequencing of FACS-sorted populations based on GFP expression shows the presence of transposon insertions at different chromosomal positions. Coverage of the insertions is uneven and sparse (Fig. 1B,D,E). In addition, there is a bias with respect to genome coordinate, with a higher insertion frequency close to the replication origin and a lower insertion frequency close to the terminus. The distributions of insertion frequencies in the parental populations of the P1-short and P1-long promoter insertions were different, but showed the same qualitative features as a function of genome coordinate (Supplementary Figure S2). Populations derived from the high or medium GFP expression populations show a bias for insertions closer to the origin of replication (Supplementary Figure S2).

We tested a possible role of gene dosage in the origin-to-terminus bias of insertion frequency. The samples are in early log phase in LB medium at 37*°C* when they are exposed to the transposon. There is therefore a higher number of copies of the chromosome close to the origin than to the terminus. Estimating the dosage from the Cooper-Helmstetter model [33], and assuming an insertion rate proportional to the dosage, we computed the expected insertion bias, keeping into account the population age-structure (see SI text).

Fig. 1B shows that the dosage estimated theoretically agrees very well with whole-genome sequencing of genome copy number, but is not sufficient to explain the stronger origin-terminus bias of the insertions. We also verified that this bias was not due to the insertions with top 10% coverage and to the insertions on ribosomal genes. The additional bias may be due to additional factors such as DNA supercoiling or biased binding of nucleoid proteins and differences in nucleoid compaction [34, 35, 36, 37]. Additionally, the density of insertions shows a slight left-right asymmetry with respect to the origin, visible when the sliding average of insertions is compared with the prediction from dosage (Fig. 1B).

### 3.4 H-NS binding sites are enriched at insertions positions

In order to better characterize the genomic positions of the insertions, we investigated the statistical tendencies for localization of insertions using the gene lists from the NuST database [32]. This database contains a large panel of published gene sets measuring several genomic properties such as binding of nucleoid-associated proteins, including several H-NS data sets (Supplementary Table 1). To score for significance, we compared the co-occurrences of insertions and genes with 5000 realizations of a shuffling null model (see Supplementary Methods for details). Note that the null model subtracts the empirical sliding average of insertions, and not the dosage, thus the results are net of the overall enrichment around the origin. The analysis was applied to the population-sequencing data for the insertion sites in both the parental populations as well as in the ones that were sorted for gene-expression levels.

This analysis, summarized in Fig. 2A, shows that H-NS binding is the main property associated with any insertions (even before any sorting by gene expression is performed), and is shared with putative horizontal transfers detected from sequence properties (among which AT-richness [38]). Indeed, the full list of insertion sites is enriched in H-NS target genes regardless of the expression level (Supplementary Table 2). We also found an enrichment on H-NS binding sites in the surroundings (10kb regions) of the insertions compared to random sites (Supplementary Fig. S2).

**Figure 2:**
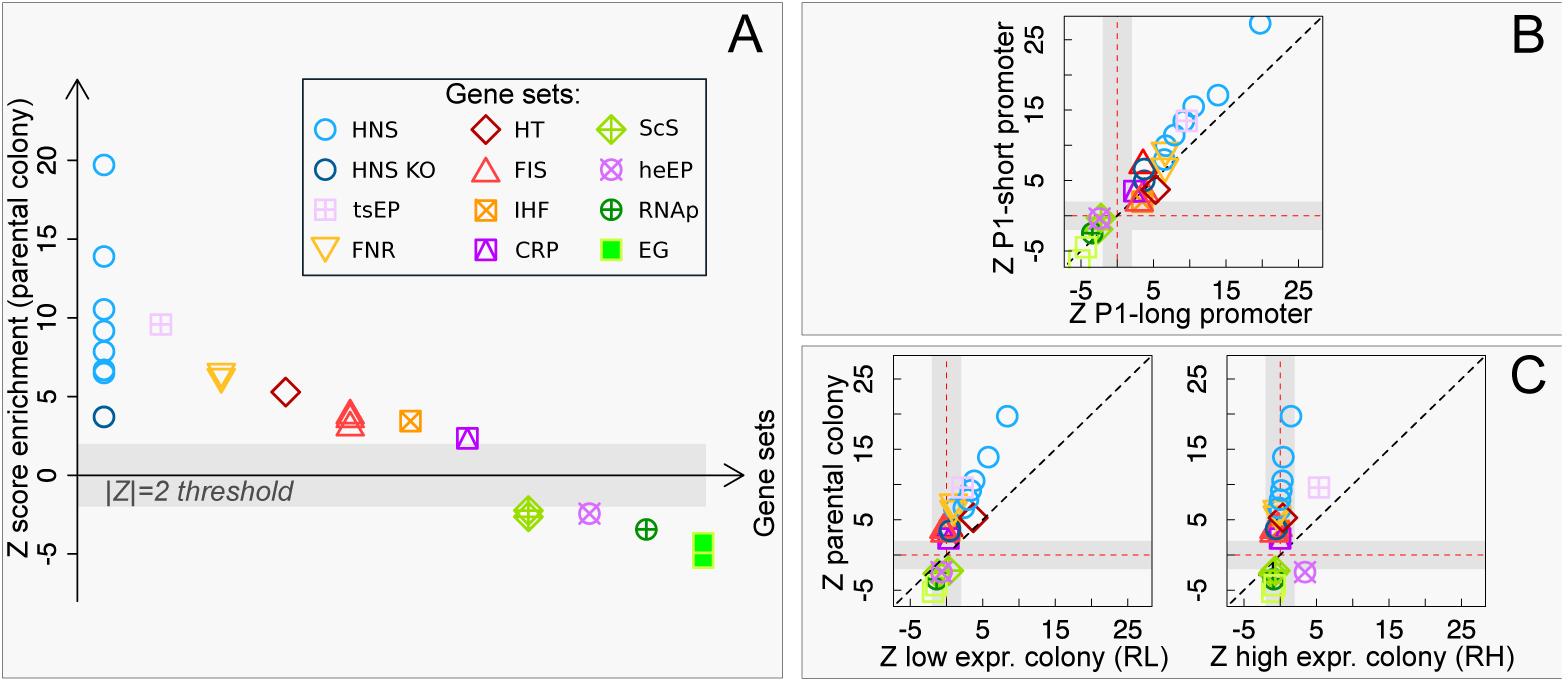
Enrichment of insertions for H-NS and other global regulators. A. Z-score of enrichment tests for different gene lists (See SI Table 1 for a full legend). H-NS binding sites (from ChIP-seq and ChIp-ChIP data) and H-NS perturbations experiments (from ref. [42]) are highly enriched (circles), indicating a strong positive association of insertions to H-NS binding regions starting from the parental colony. Other global nucleoid regulators (FNR, Fis, IHF, CRP, see legend), and a list of horizontal transfer genes (HT, see legend) also show positive association, lists of essential genes (filled squares) show strong negative enrichment. B. Comparison of the two different promoter tested (with and without Fis and H-NS binding sites) shows a similar behavior. C. Comparison of parental and sorted populations (see Fig. 1CDE) shows that H-NS association maintains a strong significance in the low-expression population, and loses significance in the high-expression population, where FNR sites remain highly enriched.

The positive local association of H-NS binding sites with insertion sites is in agreement with a common preference for AT-rich regions. As previously mentioned, genomic insertions have a reported bias for AT-rich regions [3, 1, 24], and AT-rich regions are also the preferred binding targets for H-NS [10, 19, 9, 22]. We looked for a direct correlation between insertions and AT-rich regions. In order to do this we compared the distribution of AT-bias in the sequences surrounding insertions with a random sample of the background sequences of the genome. A one-tailed Kolmogorov-Smirnov test (p-value < 10^*-*16^) suggests that there is a significant difference between the two distributions, with the sequences surrounding the insertions being richer in AT than the background ones (Supplementary Fig. S2). The role of H-NS has been proposed to inhibiting insertions in addition to repressing events of spurious transcription [12, 39, 40, 18]. Our results lead us to conclude that, in the tested conditions, at these fast growth rates, H-NS does not appear to inhibit physical events of transposon insertion efficiently. Fig. 2B shows that the results are consistent across the two promoters (P1-short and P1-long) used here (see also Supplementary Table 5), which have different sensitivity to Fis and H-NS binding and thus different intrinsic activity (also depending on growth conditions).

### 3.5 Other global regulators are enriched at insertions positions

Other gene sets that share enrichment for any insertions are targets of global regulators Fis (which alters the nucleoid state to aid transcription in exponential growth) and FNR (which alters the distribution of RNA polymerase in response to oxygen starvation). This could be related to AT-richness bias of the binding site of these proteins or to high transcriptional activity (and thus accessibility for insertions) of these genes [39, 40]. It is reasonable to expect that Fis targets are more active in LB medium. However, we found that transcriptionally active RNAP binding regions (measured by ChIP-chip during rapid growth [41]) are *under-represented* for insertions, which suggests a negative interaction between RNAP binding or transcriptional activity and insertion frequency (Supplementary Table 3).

Finally, a milder but significant over-representation for insertions was found for CRP and IHF targets, genes that are sensitive to supercoiling perturbations in an H-NS knockout background, and genes with trans-membrane domains (Supplementary Table 4). Conversely, essential genes are under-represented for insertions, as expected (Supplementary Table 3).

### 3.6 H-NS binding is the sole over-represented signal in low-expression clonal populations

Finally, we can compare the parental population with the ones sorted for gene expression levels (Fig. 2C and Supplementary Table 6). The comparison of the insertion sites of the high and low expressing populations from the P1-long promoter strains shows that the low expressing population is found preferentially within H-NS binding regions, while the high expressing population is not. Indeed, the low-expression (RL) population maintains a similar association as the parental population with H-NS and no other binding protein. Conversely, the RH (high expression) populations show no association with H-NS, and only maintain some enrichment with FNR (Fig. 2C and Supplementary Table 6). Hence, despite the lack of an effect in inhibiting transposon insertion, H-NS does appear to regulate the level of gene expression of the inserted sequences.

Overall, these results point to a more complex role than is expected for H-NS in modulating genome accessibility and gene expression of recently acquired genes.

### 3.7 Flow-cytometry analysis of clonal populations shows variable noise and gene-expression properties

Each of the sorted populations was plated separately to yield individual isogenic (clonal) colonies. To gain further insight into these differences in gene expression, 658 individual clones were hand-picked from the different populations and grown in 96-well plates to measure the average fluorescence and its standard deviation by flow cytometry. From these, 90 clones where chosen to measure fluorescence and growth rate in a plate reader in different growth media.

This analysis yields the following main results (see Supplementary Figures S3,4,5)

- There is agreement between a clone’s level of fluorescence and the average level of fluorescence of its original population (Supplementary Fig. S3), as measured by both flow cytometry and the fluorimeter. Specifically, the clones from the low-expressing populations have a significantly lower average level of fluorescence.
- The magnitude of the difference between high- and low-expressing clones depends on the growth medium. The difference is greater in the faster growth medium (Supplementary Fig. S4).
- The very low-expression strains (Fig. 1E) show low expression regardless of the growth media, except for a few outliers showing out-of-trend expression (higher than expected) in the poorer medium (Supplementary Fig. S4). This is consistent with the known growth-rate dependence of H-NS activity and ribosomal operons transcription rate [43, 16, 44]. This result shows that different regions of the genome can have different growth-rate dependent properties.
- Each clonal population (with a single insertion site, as verified by sequencing) shows a distribution of fluorescence whose spread does not depend solely on the average expression level (Fig. 3A and Supplementary Fig. S5).

**Figure 3:**
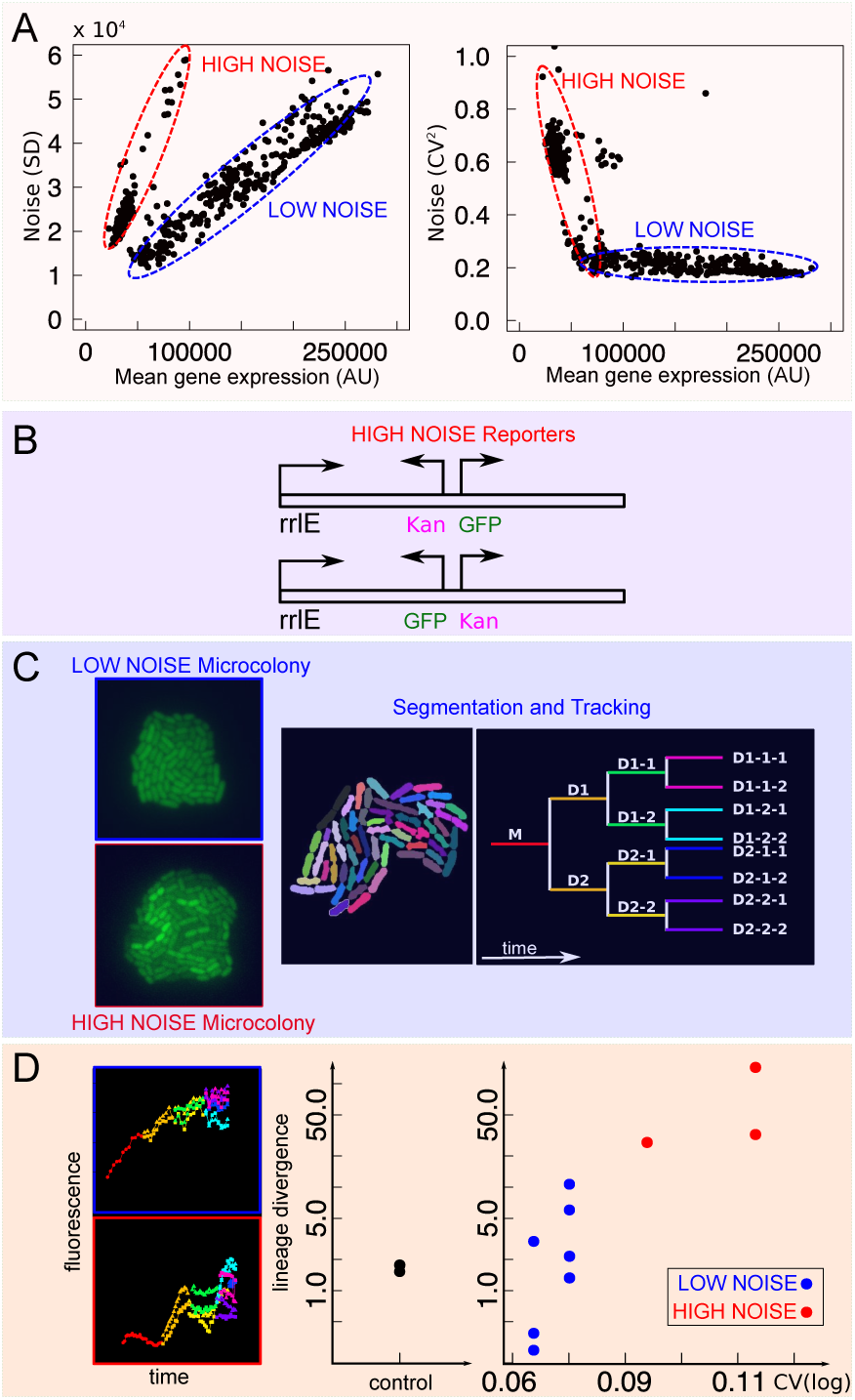
Noisy promoters emerge from transcriptional interference within the rrlE ribosomal operon. A. Characterization of noise of 658 clonal populations from flow cytometry. Left: Standard deviation vs mean GFP expression. Right: Noise (CV2) vs mean GFP expression B. All the tested “noisy” clones by sequencing typically show association with the *rrlE* ribosomal RNA operon (see Supplementary File SF1). C: Characterization of promoter noise by microcolony time-lapse growth assay. Microcolonies were grown, imaged, segmented and tracked for 3 generations. D. An example of the resulting lineage-specific gene expression data is shown on the left. The plot on the right quantifies the lineage divergence of gene expression (time average of the absolute gene expression difference) of the different clones, comparing it with the CV measured from flow cytometry (see text). High-noise clones show a large lineage divergence - indicating a possible switching behavior.

### 3.8 A set of “noisy” insertion sites

We found that in all the clones the standard deviation of gene expression is proportional to the mean, meaning that most of the noise shown by the promoter is extrinsic, regardless of expression level and insertion location (Fig. 3A). This is expected from the high level of expression of the rrnBP1 promoter [45]. However, a subset of clones show a higher level of noise where the scaling of the standard deviation with the mean follows a steeper slope. In other words, these clones show much larger gene expression variations than expected. All of these high-noise clones were very-low expression bacteria obtained from the RL (low-expression filtered) population. Hence, these clones likely explain the very-low expression sub-population (in green in Fig. 1E) in the distribution of gene expression of the RL (low-expression filtered) population. When tested for their distribution in single-cell gene expression by flow cytometry, these low-expression high-noise clones did not show bimodal distributions. Rather, they showed disperse and skewed distributions, whose range overlaps with the expression level of low-expressing clones (Supplementary Fig. S5A,B). Supplementary Fig. S5 C-G recapitulates the noise properties of all the picked clones from different rounds of selection. It is clear that high-noise clones become more frequent in successive sorting rounds where low or very-low expression cells are selected.

### 3.9 Noisy sites are associated with the insertions within ribosomal operons

To characterize the the set of clonal colonies from different populations, we performed whole-genome sequencing on 90 selected clones. The locations of all the insertions of these clones are listed in Supplementary File 1. 32 clones out of the 90 selected samples show the presence of insertion within a rRNA operon from whole-genome sequencing data. These clones were very-low expressing clones derived from RL filtered populations. In order to verify these short-read assignments of insertions sites, long-read nanopore sequencing was used on 5 of these 32 clonal samples. This analysis confirmed the presence of the transposon insertion within the 23S ribosomal RNA (rrlE) of the rrnE operon. To recapitulate these results, Fig. S6A shows the association of clones with high-noise promoters with rrlE insertions. Comparing sequenced clones with single and multiple insertions, we found that normalization of gene expression by the copy number of the promoter explained the behavior of some exceptions (Supplementary Fig. S6). Hence, we believe that the high-noise phenotype should be almost exclusively be associated with rrlE insertions.

Insertion within ribosomal operons is tolerated because *E. coli* has 7 copies of ribosomal operons. To discover the orientation of the cloned sequences with respect to the rrlE promoter position, we used the blast results of *de novo* assembled contigs with the flanks of cloned insert sequence and we compared them with the reference genome. Most of the insertions showed an opposite orientation of the inserted promoter with respect to the rrlE promoter sequence.

Separately, we checked the orientation of the inserted GFP cassette in few selected Illumina samples for which the blast results showed reasonable overlap with the genome locus. This confirmed the opposite orientation of GFP with respect to the rrlE promoter sequence in most samples. Strong promoter competition may explain the very low levels of GFP transcript production by which these clones were isolated, as well as the high variability.

We found that the trends of gene expression with growth rate were consistent with the hypothesis of competition with a strong promoter: while most of the other clones increase their expression with growth rate (in agreement with the known regulation of the rrnBP1 ribosomal promoter) the noisy clones decrease in expression with increasing growth rate, in agreement with the idea that their expression is repressed by an interference with transcription of the increasingly transcribed ribosomal operon (Supplementary Fig. S3 and S4). Additionally, the mild reduction in growth rate for insertions giving different mean GFP expression suggests that the cost associated to GFP expression is not dominant for these insertions (Supplementary Fig. S3 and S4).

### 3.10 Noisy promoters may perform switching

The previous analyses strongly indicate an association of the high-noise insertions with interference of the insertions with ribosomal operons. To gain more insight on the temporal dynamics of these high-noise inserted promoters, we measured gene expression noise in time-lapse microscopy data on growing microcolonies (Fig. 3CD). We compared the change in GFP gene expression over time of single cells from clones carrying noisy and non-noisy promoters for 3-4 generations and quantified the differences between gene expression in different lineages. The divergence between lineages was quantified as the time average of the absolute value of the gene expression difference between sister cells.

Bacteria were grown on an agar pad to form a microcolony. The time-lapse data in the formation of the microcolony was segmented to obtain the change in the average cell fluorescence as a function of time Fig. 3C. An example of gene expression of two lineages, one from a noisy clone, as measured by flow cytometry, and one from a control clone (where the cassette was inserted specifically between two converging genes, AidB and yjfN) is shown in the left panel of Fig. 3D. The right panel of Fig. 3D quantifies the divergence of gene expression along lineages for different clonal microcolonies, corresponding to clones where the promoter is inserted in different positions. Figure 3D shows that in microcolonies from high-noise clones different lineages emanating from the same single cell tend to diverge more in gene expression as time progresses than in reference or low-noise clones. This result points to the possible presence of switching behavior in the high-noise clones.

## 4 DISCUSSION

Our results directly show that the probability of DNA insertion in the *E. coli* genome by a transposon is biased, before any long-term selection may act, other than that related to overnight growth. First, there is a stronger origin-to-terminus bias than justified by gene dosage imbalance, second, insertion probability is higher in AT-rich regions of the genome. This may seem surprising, because these are regions that are preferentially bound by the H-NS protein, which has been proposed to act as a barrier to horizontal gene transfer [46, 8, 10, 6, 27, 3]. However, physical components such as differences in DNA supercoiling [47, 48, 34] and the biophysical properties of AT-rich DNA (lower melting barriers, different stacking energy, etc.) may play a role in establishing these biases. We also note that we carried out the transposon reaction in mid exponential phase cells growing in a rich medium, LB. In these conditions the concentration of H-NS is lower due to a high dilution rate [16]. However, at the faster growth rates, H-NS is known to still play a role in the repression of ribosomal promoters by binding to higher affinity sites [36, 49]. Our results suggest that there is probably not enough protein to also cover the lower affinity (nonspecific) binding to AT-rich regions, in order to inhibit transposon insertion [11]. This is in contrast to a previous study in *Vibrio cholerae* that has shown that only in the absence of H-NS there was a higher probability of insertion in AT-rich regions of the genome [12]. These results lead us to further suggest that the role of H-NS in regulating the probability of genomic insertion of horizontally acquired genes may depend on the growth conditions and on the specific strain.

Our results are consistent with the role of H-NS as a silencer of newly acquired genes. Indeed, we observe that once the full length rrnBP1 promoter cassette is inserted in the genome, its level of expression is lower if it is found near H-NS rich regions. This cassette includes a higher affinity H-NS binding site within the full length rrnBP1 promoter and an AT-rich *gfpmut2* gene sequence stabilizing the formation of H-NS dependent repressing complex. The shorter version of the promoter (P1-short), lacking the high affinity H-NS binding site, does not appear show a stable sub-population of very-low fluorescence clones (data not shown), showing that the high affinity site of the promoter is important in nucleating the repressing oligomeric structure, a question that we are still exploring. In summary, the level of expression of the clones can thus be very heterogeneous, depending on local properties of the site of insertion and the sequence of the fragment.

The cell-to-cell variability of gene expression within a given isogenic clone can vary significantly, but typically scales with the mean level of expression as expected from extrinsic noise [50, 45, 43]. This is expected from the promoter used here, rrnBP1, which is a strong promoter, resulting in a high level of expression. The change in the CV as a function of mean expression therefore remains for the most part relatively flat, corresponding to the extrinsic noise regime.

However, in some of the very low expression clones the noise varies in a way that is not expected from the known pattern of gene expression noise correlations that have been described previously [43, 45]. We therefore characterized those clones that have a higher level of gene expression noise and found that in these cases the insertion has taken place within a ribosomal operon. *E. coli* has 7 copies of ribosomal operons, therefore insertion inside one of them does not have a high cost and is not selected against, at least in the short term of this experiment. This results in interference between two transcription processes driven by very similar promoters, of similar strength. Furthermore, the initiation frequency of ribosomal promoters is high enough at fast growth rates that most of the time the operon sequence can be assumed to be covered by transcribing RNA polymerase “trains” [51]. This creates a block for RNA polymerase to bind to the promoter that is found within the operon, creating a stable “off state”. However, from time to time RNA polymerase manages to bind to the newly inserted promoter, perhaps after the DNA replication forks have erased the memory from the competing process, starting its own “train” of GFP production. Such transcriptional interference is a well-known phenomenon [52]. Our result on promoter noise also suggests that in the tightly packed bacterial genomes, transcription interference with newly inserted genes might be a natural source of innovations in terms of gene expression noise on evolutionary time scales, as previously speculated for eukaryotes [53].

Altogether, our findings support the following evolutionary scenario. When a novel gene enters the genome, it is more likely found in a region that is controlled by H-NS, for reasons that most likely have nothing to do with fitness, but have to do with the physico-chemical properties of the DNA in AT-rich regions. However, the wide range of expression levels that we find show that the gene is not necessarily immediately silenced. Rather, the different insertion positions allow it to sample a wide range of expression levels (including silencing), at (initially) equal promoter strength, while interacting from the start with the cell’s housekeeping physiology. We believe that this inherent bet-hedging and exploratory stage may be a key ingredient of genome plasticity, and is underestimated in our current narrative of the process of horizontal transfer, which is centered on the average outcome, and establishes a strict time hierarchy between stages where an exogenous gene is first silenced and then reactivated.

## 5 ACKNOWLEDGEMENTS

This work was made possible by the support from the IFCPAR/CEFIPRA (Indo-French Centre for the Promotion of Advanced Research) Grant 5103-3.

