## Supplementary for "Early fate of Exogenous promoters in *E. coli*"

Supplementary materials for Yousuf *et al.*
"Early fate of exogenous promoters in *E. coli*"

### Supplementary Methods

#### Estimate of the expected gene-dosage effect in transposition.

This section discusses the model used to understand a null trend in the insertions coming from gene dosage (1; 2). The samples are grown in rich medium and when they are exposed to the transposon, there are a higher number of copies of the chromosome close to the origin than close to the terminus (Figure 1 in the main text).

We proceed to estimate the dosage theoretically. We assume that the $C + D$ periods last 40 + 20 minutes, since we are dealing with fast growth in LB medium, and that the doubling time $\tau$ is around 25 minutes (which is the most conservative estimate in terms of dosage bias). Since at time $C + D$ ($\approx 60$ minutes) the cell divides, a time lag $B$ before initiation is necessary to make the total replication time $B + C + D$ an integer multiple of the doubling time $\tau$, âĂIJsynchronizingâĂİ DNA replication and cell division. Thus, defining $n = \mathrm{Int}\frac{C+D}{\tau}$ as the integer number of times that $\tau$ divides $C + D$, the following relation has to be satisfied:

$$B + C + D = (n + 1)\tau \tag{S1}$$

More generally, we can consider a gene at a chromosomal position defined by its normalized distance from Ori, i.e. *l* $= 0$ represents a gene in Ori and *l* $= 1$ in Ter. The copy number of this gene, $g$ , changes during the cell cycle following:

$$g(t) := \begin{cases} 2^{n'} & \text{if } 0 < t < (n'+1)\tau - (c(1-l) + D) \\ 2^{n'+1} & \text{if } (n'+1)\tau - (c(1-l) + D) < t < \tau \end{cases} \tag{S2}$$

where $n' = \mathrm{Int}\left[\frac{C(1-l)+D}{\tau}\right]$.

Next, the population age structure must be taken into account when evaluating the average gene dosage in a cell population. For ideal balanced exponential growth with rate $\mu$ this distribution is given by $a(t, \mu) = 2\ln(2)\mu 2^{-\mu t}$.

Thus, the population average is:

$$\langle g(t) \rangle_{population} = \int_0^\tau a(t,\mu) g(t) dt = 2^{\mu[C(1-l)+D]}. \tag{S3}$$

We normalize the basepair position from 0 to 1 where 0 is origin and 1 is terminus.

In order to find a correct estimate of the mean dosage of insertions, we also have to introduce a model for insertion kinetics. The simplest model relies on the assumption of constant insertion rate. We hypothesize that during a time interval $t$ a cell population that grows exponentially with division time $\langle \tau \rangle$ is exposed to a transposon which inserts its sequence at a constant rate $r$ per time and genome coordinate unit. The expected number of insertions under this assumption can be obtained by the cumulative distribution function. The mean copy number of each coordinate in a cell population is $c(x)$ and we can assume that there is a Poisson process with the same rate in each genome position, so that the dependency of the rate from position $x$ is only due to dosage, $r(x) = c(x)S$. In this case we get,

$$P(x) = 1 - e^{r(x)t} \simeq r(x)T \simeq c(x)ST = Ac(x) \ , \tag{S4}$$

where we have linearized the expression for small $t$. This expression gives an estimate for the insertion probability , with one free parameter $A = ST$, and depending on the measured parameters $C,\ D,\ \mu$. The trend of normalized position and the dosage estimate for insertions is shown in Fig.1B in the main text. This plot shows that this estimate does not correspond well quantitatively to the experimental coverage data.

We verified that a model with time-dependent insertion rate, i.e. where the insertion rate $r$ *increases* with time $t$, $r(x,t)$, can fit the data, using insertion rate growing as a power law in time. However, there is no empirical motivation to assume such cooperative behavior in insertions that occur *in different cells*.

Our final consideration concerns the effect of exponential growth of the different inserted clones. Let $f_i$ be the fraction of insertions in site $i$ and $f_j$ the fraction of insertions in site $j$. If $N$ is the total number of insertions we will have, at time $\tau$, $N * f_i e^{\alpha\tau}$ insertions in site $i$ and $N * f_j e^{\alpha\tau}$ insertions in site $j$. Then, the total number of insertions will be $Ne^{\alpha\tau}$.

Let $f_{i,j}(\tau)$ be the ratio of the numbers of insertions in site $i$ and $j$. Then,

$$f_{i,j}(\tau) = \frac{N * f_{i,j} e^{\alpha\tau}}{Ne^{\alpha\tau}} = f_{i,j} \tag{S5}$$

We have found $f_{i,j}(\tau) = f_{i,j}$, showing that, for all those insertions not affecting the growth rate, exponential propagation of insertions in progeny does not create a bias in the population. If instead an insertion affects the growth rate of the corresponding clone its presence will be clearly biased in the population.

#### Null model for insertion enrichment.

This section describes the null model used to score enrichment of insertions in particular regions defined by gene lists. Since the thoeretical prediction scores poorly (and even a fit with a time-dependent insertion rate model does not capture qunaitativley the shape of the dosage peak around the origin), in the analysis for scoring overlap between insertions and gene lists we decided to use a more conservative way to subtract the dosage, i.e. directly to consider a sliding-window average of the data as background insertion probability. Specifically, we used a window size of 3000 bp, which is the smallest window size that gave a smooth curve with few local features.

In order to investigate statistical tendencies for transposon insertions to be associated to specific chromosomal contexts, we need a suitable null model. This is simple to define using gene lists, as randomizations of the empirical lists having fixed number of genes. In particular, for each tested gene list we simulate 5000 different stochastic realizations.

The definition of enrichment needs a quantity representative of the tendency of finding the insertion fixed in specific chromosomal contexts. We evaluate a discrete integral considering the coverage corresponding to a portion of genome with "switch on" genes. We call this quantity "coverage integral". It corresponds to the total overlap of the coverage with the genes of a specific list.

We have evaluated the Z-score as significance score for over/under-representation of the overlap between coverage and target gene lists. The Z-score is defined as $Z = \frac{X - \mu}{\sigma}$ where $X$ is the empirical value and $\mu$ and $\sigma$ are the mean and the variance relative to a particular sample.

##### DNA primers used:

To verify the insert on the Tn5 construction vector:
Forward Primer : AATTCTACGAGCGTCGTCGCAGACATGATC
Reverse Primer : GGTCTCAGATGGTAGACGCAGGAAGAGACGAGACAG

Primers used for transposon sequencing experiments (* indicates a phosphorothioate group:
Index primer: AAGAGCGGTTCAGCAGGAATGCCGAGACCGATCTC

3âĂŹ Tnspecific primer:
AATGATACGGCGACCACCGAGATCTACACCCAATATGCGAGAACACCCGAGAAAATTCATCG

3âĂŹ Sequencing primer: CCCGAGAAAATTCATCGATGATGGTTGAGATGTGTA

Illumina Read 1: ACACTCTTTCCCTACACGACGCTCTTCCGATCT
Illumina Read 2: CGGTCTCGGCATTCCTGCTGAACCGCTCTTCCGATCT

QPCR2.1 : CAAGCAGAAGACGGCATACGA
qPCR2.2 : AATGATACGGCGACCACCGAG

Adapters:
SplA5_top G*AGATCGGTCTCGGCATTCCTGCTGAACCGCTCTTCCGATC*T
SplA5_bottom /5Phos/G*ATCGGAAGAGCGGTTCAGCAGGttttttttttcaaaaaaa*a
SplAP5.2 C*AAGCAGAAGACGGCATACGAGATAAACATCGGAGATCGGTCTCGGCATTC*C
SplAP5.5 C*AAGCAGAAGACGGCATACGAGATACCACTGTGAGATCGGTCTCGGCATTC*C
SplAP5.6 C*AAGCAGAAGACGGCATACGAGATACATTGGCGAGATCGGTCTCGGCATTC*C
SplAP5.12 C*AAGCAGAAGACGGCATACGAGATAGTACAAGGAGATCGGTCTCGGCATTC*C

#### Supplementary Figures

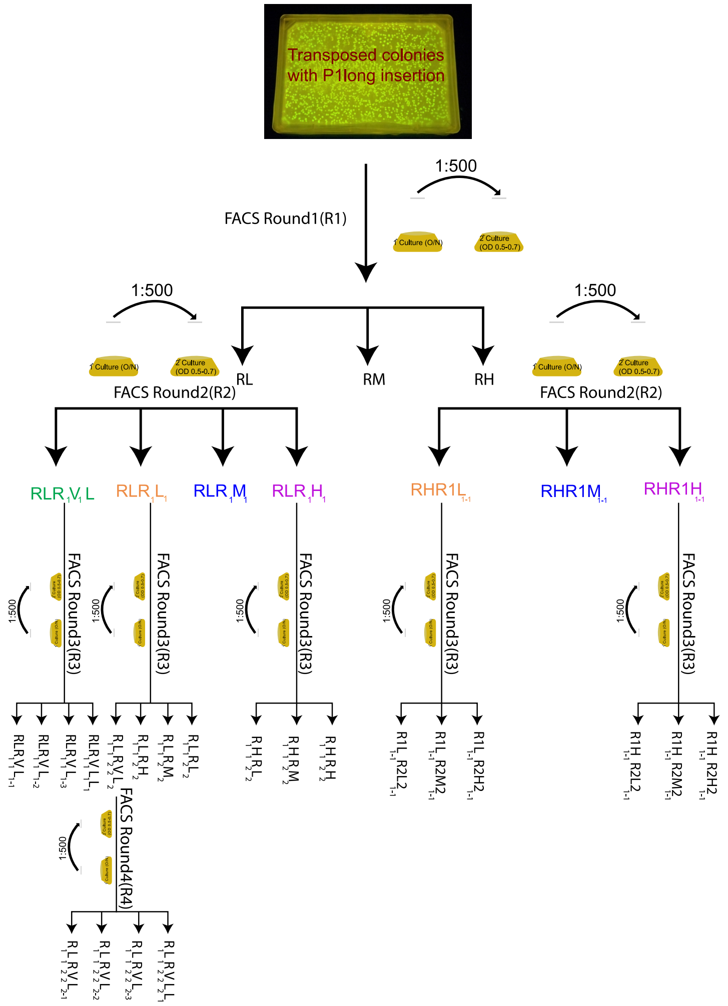


Figure S1: **Full illustration of the sorting pipeline starting from the parental population and the nomenclature used in our experiment.** The parental population (the cells that underwent transposition) was grown overnight in M9 medium with 0.5% Glucose + 0.2% CAA ("fast growth" medium) and was diluted to 1:500 to grow in secondary culture with the same medium the following day until the OD reached 0.5 âĂŞ 0.6. These cultures underwent first round of FACS to yield three population by the level of GFP expression (Low (RL), Medium (RM) and High (RH)). Subsequently the Low (RL) and High (RH) underwent the second round of FACS, the former (RL) yielded 4 population, RLR1V1L1 (Very low), RLR1L1 (Low), RLR1M1 (Medium) and RLR1H1 (High) while the latter (RH) yielded three population, RHR1L1-1 (Low), RHR1M1-1 (Medium) and RHR1H1-1 (High). From the second round of FACS, three populations, RLR1V1L1 (Very low), RLR1L1 (Low), and RLR1H1 (High) were selected from the low expressing population while 2 populations, RHR1L1-1 (Low) and RHR1H1-1 (High) from the high expressing population were selected for the third round of FACS. RLR1V1L1 (Very low) gave rise to 4 different populations, RLR1V1L1L1, RLR1V1L1-1, RLR1V1L1-2, RLR1V1L1-3 and RLR1L1 (Low) also gave rise to 4 different population, R1L1R2V2L2, R1L1R2L2, R1L1R2M2, R1L1R2H2 while RLR1H1 (High) gave rise to 3 different populations, R1H1R2L2, R1H1R2M2 and R1H1R2H2. From the high expressing populations, RHR1L1-1 and RHR1H1-1 gave rise to R1L1-1R2L21-1, R1L1-1R2M21-1, R1L1-1R2H21-1, and R1H1-1R2L21-1, R1H1-1R2M21-1, R1H1-1R2H21-1, respectively. Only the very low expressing population, R1L1R2V2L2 from the third round of FACS gave rise to 4 different very low expressing populations, R1L1R2V2L2-1, R1L1R2V2L2-2, R1L1R2V2L2-3, R1L1R2V2L2-4.

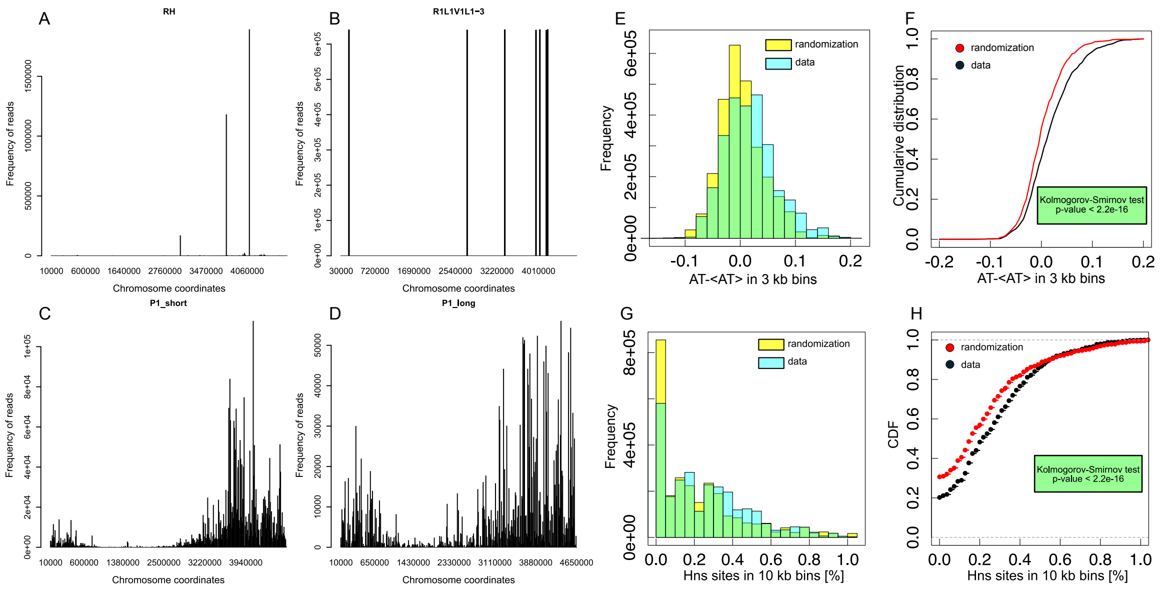


Figure S2: **Insertion frequency plots and AT bias.** A): Highly expressing sorted (RH) population shown in Figure 1D shown in linear scale. B): Insertions in low expressing sorted population (RL) shown in Figure 1E, shown in linear scale, transposon insertions in a rRNA operon are visible. C) and D): Insertion frequency plots for parental populations of P1-long and P1-short promoters respectively. E) and F) Insertions have a significant positive bias in AT-richness compared to the background. E): Histograms of deviations from average in AT-richness ($\%AT- < \%AT >$) in the sequences surrounding insertions from the experimental data and from a randomized sample of the background sequences of the genome. $\%AT$ corresponds to a local average in 3Kb intervals around each insertion; $< \%AT >$ refers to the average on the entire genome. F): comparison of the cumulative distributions and results of the Kolmogorov-Smirnov test (in both cases the p-value is smaller than the smallest represented by the ks.test R script). G) and H) Insertions have a positive bias for H-NS binding. The plots report the distribution of the fraction of the 10kb region around an insertion covered by H-NS binding sites (ChIPseq data from ref. (3)), compared to the same quantity for randomized insertion sites.

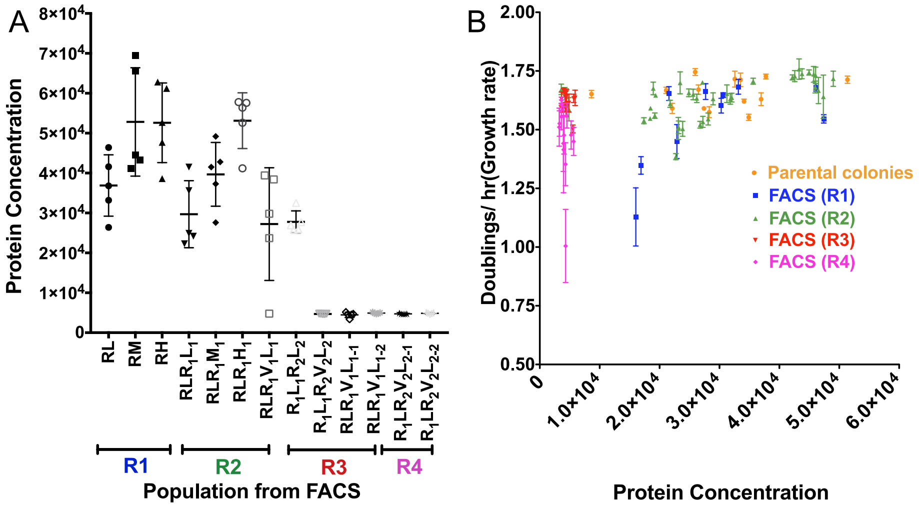


Figure S3: **Expression levels of clones from sorted populations measured by fluorimetry, and growth rate.** (A) The fluorimeter expression data are in agreement with the FACS population histograms. GFP expression (total fluorescence over OD) of 5 individual strains randomly chosen from each of the different rounds of FACS sorted population. All the strains were grown in rich medium (0.5% Glucose + 0.2% CAA). Error bars represent standard error of the mean (SEM). R1, R2, R3, R4 denote the first, second, third and fourth round of FACS. Expression is low for very low expressing population (VL in R3 and in R4) and significantly different from the other sorted populations ($P < 0.001$, One-way ANOVA, Tukey's multiple comparison test comparing R1, R2 with R3 and R4). The fluorimeter expression data are in agreement with the FACS population histograms. (B) Scatter plot of expression levels and population growth rates for clones from the different rounds.

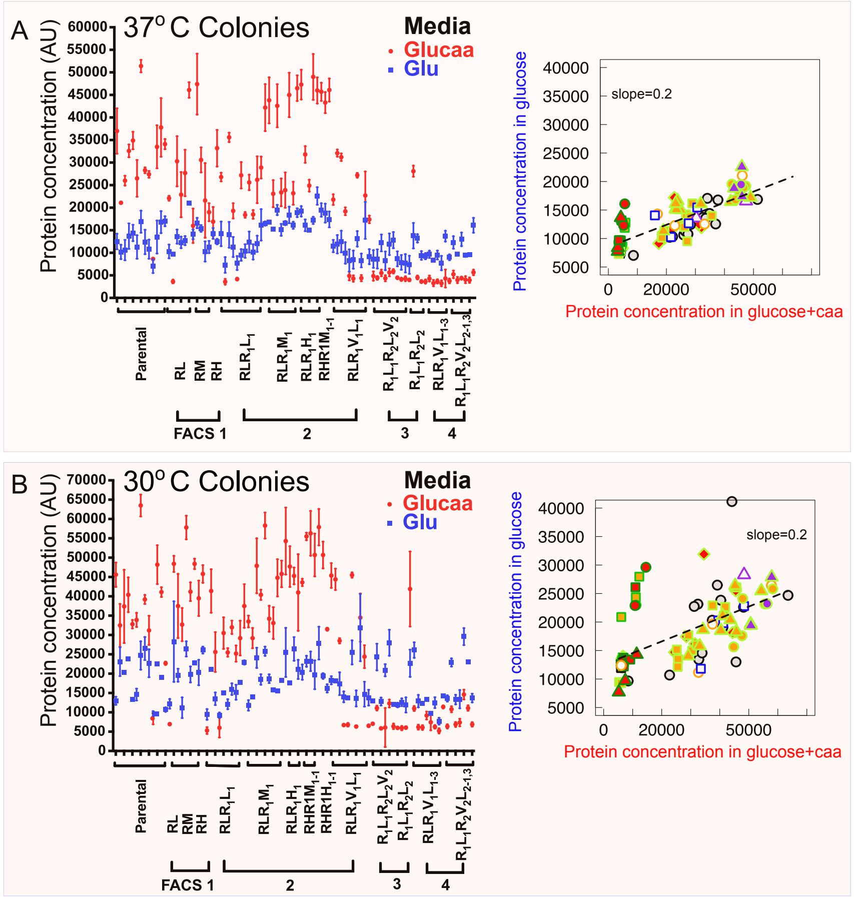


Figure S4: **Insertions at the ribosomal regions are highly repressed in different growth conditions.** The expression levels of these colonies are lower when they are grown in a fast-growth medium (M9 + 0.5% Glucose + 0.2% CAA) compared to the slower-growth medium (M9 + 0.5% Glucose) irrespective of the temperature grown (37$^{o}$C and 30$^{o}$C). Panel A) reports the gene expression of 90 clones from different rounds of FACS grown in different media (fast growth, Glucose + CAA, compared to the slow growth medium, Glucose at 37$^{o}$C). Error bars represents standard deviation (SD), n = 3. Panel B) represents the comparison of colonies grown in different media (fast vs slow growth medium) at 30$^{o}$C, error bars represent SD, n = 2. The scatter plots on the right side of each panel compare gene expression of the same clone in the two growth conditions (color/symbol codes defined in Supplementary Fig. 5). Note that while most of the other clones increase in expression in the faster growth medium (in agreement with a ribosomal promoter) the very-low expression (VL) ones decrease with the faster growth rate, in agreement with the idea that their expression is decreased by the activity of a ribosomal promoter. The comparison between the two temperatures shows that there is a subset of clones that increase in expression in the slower growth Glucose medium at low temperature, spanning different sorted populations. Note that thereare two growth rate dependence results, the first is the fact that we grew the cells in two growth media and we observed an increase in gene expression in the richer medium, as expected from the rrnBP1 promoter, the second is that within a given growth medium different clones grew at different rates (Fig. S3B).

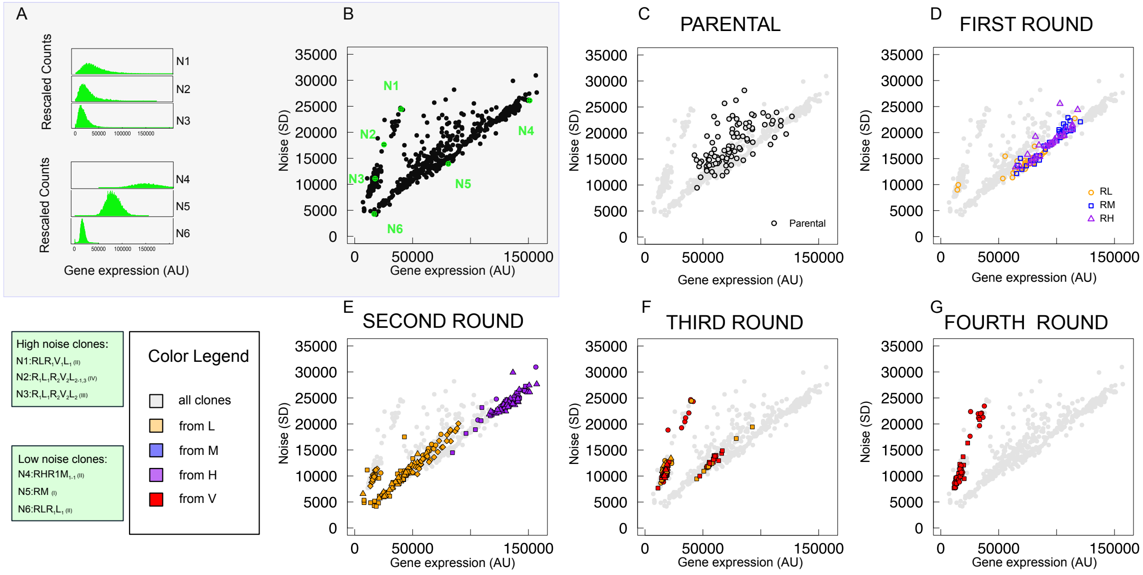


Figure S5: **Gene expression levels and noise of clonal populations from FACS.** The figure recapitulates the main results of the selected clonal populations (hence carrying the same insertions). A) and B) N1-N6 are six representative clones chosen from all the measured ones. Panel B (the same plot shown in Figure 3A) shows their gene expression mean level and noise (SD), while panel A compares the histograms of their gene expresion levels. Panels C-G use the same plot to show the noise properties of the clones coming from different rounds of FACS sorting.

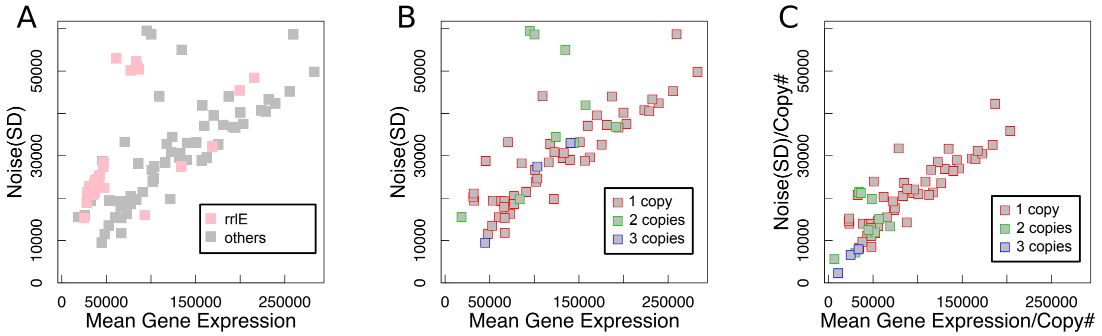


Figure S6: **Gene expression noise categories of sequenced clones with varying insertion copy number.** A) Mean vs SD plots of rrlE vs non-rrlE insertions. Noisy promoters mostly associate with rrlE insertions in sequenced clones. B) Mean vs SD plots of non-rrlE sequenced clonal populations (a subset of the data in Supplementary Fig. S5), where the points are colored by number of insertions. C) Different plot of the same data, where gene expression normalized per copy number. Removal of the effect of copy number fold change in the expression levels reconducts many rrlE outliers to the cluster of low-noise promoters.

#### Supplementary Tables

| Genes linked to specific phenotypes (essential genes and genes responsive to external conditions) | | | | | | | | | Mutated genes | | |
|---|---|---|---|---|---|---|---|---|---|---|---|
| Multi-stress responsive genes (from Nichols et al 2011) | Conditionally essential genes due to auxotrophy (from Nichols et al 2011) | Conditionally essential genes in rich media (from Nichols et al 2011) | Genes whose expression is affected by growth conditions (from Regulon DB) | Genes whose expression is affected by the carbon source (from RegulonDB) | List of E. coli essential genes (from Baba et al 2006) | List of E. coli essential genes (from Gerdes et al 2003) | | | List of mutated genes in E. coli long-term evolution experiment through 20,000 generations (Barrick et al 2009) | List of mutated genes in E. coli long-term evolution experiment through 40,000 generations (Barrick et al 2009) | |
| Genes next to binding sites of a regulator from binding data (ChIP-chip, ChIP-seq, etc) | | | | | | | | | | | |
| Putative RNA-polymerase target genes during rapid growth from ChIP-chip experiments (from Grainger et al 2005) | Genes overlapping with heEPODs (highly expressed extended protein occupancy domains) (from Vora et al 2009) | Genes overlapping with tsEPODs (transcriptionally silent extended protein occupancy domains) (from Vora et al 2009) | Putative CRP target genes from ChIP-chip experiments (from Grainger et al 2005) | Putative FIS target genes from ChIP-chip experiments (from Cho et al 2006) | Putative FIS target genes from ChIP-seq experiments (Kahramanoglu et al 2010) | Putative FIS target genes in early-exponential phase from ChIP-seq experiments (from Kahramanoglu et al 2010) | Putative FIS target genes in mid-exponential phase from ChIP-seq experiments (from Kahramanoglu et al 2010) | Putative FIS target genes in mid-log phase from ChIP-chip experiments (from Grainger et al 2006) | Putative FNR target genes in stationary phase from ChIP-chip experiments (from Grainger et al 2007) | Putative FNR target genes in midlog phase in the presence of oxygen from ChIP-chip experiments (from Grainger et al 2007) | Putative FNR target genes in midlog phase from ChIP-chip experiments (Grainger et al. 2007) |
| Putative H-NS target genes from ChIP-chip experiments (from Oshima et al 2006) | Putative H-NS target genes from ChIP-seq experiments (Kahramanoglu et al 2010) | Putative H-NS target genes in early-exponential phase from ChIP-seq experiments (from Kahramanoglu et al 2010) | Putative H-NS target genes in mid-exponential phase from ChIP-seq experiments (from Kahramanoglu et al 2010) | Putative H-NS target genes in mid-log phase from ChIP-chip experiments (from Grainger et al 2006) | Putative H-NS target genes in stationary phase from ChIP-seq experiments (from Kahramanoglu et al 2010) | Putative H-NS target genes in transition-to-stationary phase from ChIP-seq experiments (from Kahramanoglu et al 2010) | Putative IHF target genes in mid-log phase from ChIP-chip experiments (from Grainger et al 2006) | | | | |
| Genes sensitive to nucleoid perturbations from transcriptomics experiments | | | | | | | | | | | |
| FIS knockout sensitive genes in all stages of growth (from Bradley et al 2007) | FIS knockout sensitive genes in the early-exponential growth phase (from Bradley et al 2007) | FIS knockout sensitive genes in the late-exponential growth phase (from Bradley et al 2007) | FIS knockout sensitive genes in the mid-exponential growth phase (from Bradley et al 2007) | FIS knockout sensitive genes in the stationary phase (from Bradley et al 2007) | FIS-knockout sensitive genes under negative-supercoiling (from Blot et al 2006) | FIS-knockout sensitive genes under positive-supercoiling (from Blot et al 2006) | H-NS knockout sensitive genes under negative-supercoiling (from Blot et al 2006) | H-NS knockout sensitive genes under positive-supercoiling (from Blot et al 2006) | Supercoiling sensitive genes under FIS knockout (from Blot et al 2006) | Supercoiling sensitive genes (from Blot et al 2006) | Supercoiling sensitive genes under H-NS knockout (from Blot et al 2006) |
| Genes with specific annotations | | | | Target genes of a regulator (according to external databases) | | | | | | | |
| Putative horizontally transferred genes (from HGT-DB) | Genes related to small RNAs (from Regulon DB) | Genes whose protein products have transmembrane domains (from Ensembl Bacteria) | Genes with paralogs (from Ensembl Bacteria) | Genes regulated by small RNAs (from Regulon DB) | Genes regulated by the sigma factor 24 (from Regulon DB) | Genes regulated by the sigma factor 28 (from Regulon DB) | Genes regulated by the sigma factor 32 (from Regulon DB) | Genes regulated by the sigma factor 38 (from Regulon DB) | Genes regulated by the sigma factor 54 (from Regulon DB) | Genes regulated by the sigma factor 70 (from Regulon DB) | |

Table S1: **Gene lists used in the enrichment analysis, grouped by category.** Lists were divided according to the types of biological data and the experimental techniques. See ref. (4) for details. The symbols are the ones used in Fig. 2 in the main text.

| GENE SET | Z-SCORE (p1 long) | Z-SCORE (p1 short) | Z-SCORE (p1 short) |
|---|---|---|---|
| Putative H-NS target genes from ChIP-seq experiments (Kahramanoglou et al 2010) | 19,7 | 27.3 | 27,3 |
| Putative H-NS target genes from ChIP-chip experiments (from Oshima et al 2006) | 13,9 | 17,1 | 17,1 |
| Putative H-NS target genes in early-exponential phase from ChIP-seq experiments (from Kahramanoglou et al 2010) | 10,5 | 15,5 | 15,7 |
| Genes overlapping with tsEPODs (transcriptionally silent extended protein occupancy domains) (from Vora et al 2009) | 9,5 | 13,5 | 13,6 |
| Putative H-NS target genes in mid-exponential phase from ChIP-seq experiments (from Kahramanoglou et al 2010) | 9,2 | 13,4 | 13 |
| Putative H-NS target genes in transition-to-stationary phase from ChIP-seq experiments (from Kahramanoglou et al 2010) | 7,9 | 11,4 | 11,5 |
| FNR target genes in midlog phase in the presence of oxygen from ChIP-chip experiments (from Grainger et al 2007) | 6,7 | 7,2 | 7,7 |
| Putative H-NS target genes in stationary phase from ChIP-seq experiments (from Kahramanoglou et al 2010) | 6,6 | 10,8 | 10 |
| Putative FNR target genes in midlog phase from ChIP-chip experiments (Grainger et al. 2007) | 6,5 | 8,9 | 9 |
| Putative FNR target genes in midlog phase in the presence of oxygen from ChIP-chip experiments (from Grainger et al 2007) | 6,4 | 7,2 | 7,2 |
| Putative horizontally transferred genes (from HGT-DB) | 5,3 | 4,1 | 4,1 |
| Putative FIS target genes from ChIP-seq experiments (Kahramanoglou et al 2010) | 3,8 | 7,1 | 7 |
| H-NS knockout sensitive genes under negative-supercoiling (from Blot et al 2006) | 3,7 | 6,6 | 6,7 |

Table S2: **Groups of genes regarding HNS or horizontally transferred genes result over-represented.** The table reports the gene sets that result over-represented (Z-score $> +5$ in at least one experiment).

| GENE SET | Z-SCORE (p1 long) | Z-SCORE (p1 short) | Z-SCORE (p1 short) |
|---|---|---|---|
| List of E. coli essential genes (from Gerdes et al 2003) | -5,2 | -6,3 | -6,2 |
| List of E. coli essential genes (from Baba et al 2006) | -4,4 | -4,5 | -4,4 |
| Putative RNA-polymerase target genes during rapid growth from ChIP-chip experiments (from Grainger et al 2005 | -3,4 | -2,5 | -2,7 |
| Supercoiling sensitive genes (from Blot et al 2006) | -2,6 | -2 | -1,8 |
| Supercoiling sensitive genes under FIS knockout (from Blot et al 2006) | -2,4 | -0,3 | -0,2 |
| Genes overlapping with heEPODs (highly expressed extended protein occupancy domains) (from Vora et al 2009) | -2,2 | -0,4 | -0,5 |

Table S3: **Groups of essential genes result under-represented.** The table reports the gene sets that result under-represented (Z score $< -3$ in at least one insertion experiment).

| GENE SET | Z-SCORE (p1 long) | Z-SCORE (p1 short) | Z-SCORE (p1 short) |
|---|---|---|---|
| Putative FIS target genes in mid-log phase from ChIP-chip experiments (from Grainger et al 2006) | 3,8 | 3,2 | 3 |
| H-NS knockout sensitive genes under positive-supercoiling(from Blot et al 2006) | 3,7 | 4,1 | 4,9 |
| Putative IHF target genes in mid-log phase from ChIP-chip experiments (from Grainger et al 2006) | 3,6 | 2 | 2,1 |
| Putative FIS target genes from ChIP-chip experiments (from Cho et al 2006) | 3,5 | 3,1 | 2,2 |
| Putative CRP target genes from ChIP-chip experiments (from Grainger et al 2005) | 2,4 | 3,5 | 3,2 |

Table S4: **Groups of Fis,IHF,CRP target genes result over-represented considering a lower limit** The table reports the gene sets that result over-represented (Z score $> 3$ and $< 5$ in at least one insertion experiment).

|  | P1 long RL | P1 long RH | P1 short 1 | P1 short 2 |
|---|---|---|---|---|
| P1 long | 0.86 | 0.50 | 0.97 | 0.97 |
| P1 long RL |  | 0.60 | 0.83 | 0.84 |
| P1 long RH |  |  | 0.51 | 0.50 |
| P1 short 1 |  |  |  | 0.99 |

Table S5: **A correlation exists between Z scores from different strains** Correlation between Z scores from different stains is represented by the Pearson correlation coefficient. P1 long is the parental colony, then we have the first round high expression (RH) and low-expression (RL) sorted populations. Different colors correspond to different correlation levels.

| POPULATION | GENE SET | Z-SCORE |
|---|---|---|
| RL | Putative H-NS target genes from ChIP-seq experiments (Kahramanoglou et al 2010) | 8,4 |
|  | Putative H-NS target genes from ChIP-chip experiments (from Oshima et al 2006) | 5,8 |
|  | Putative H-NS target genes in early-exponential phase from ChIP-seq experiments (from Kahramanoglou et al 2010) | 3,8 |
|  | Putative horizontally transferred genes (from HGT-DB) | 3,7 |
|  | Putative H-NS target genes in mid-exponential phase from ChIP-seq experiments (from Kahramanoglou et al 2010) | 3,4 |
|  | Putative H-NS target genes in transition-to-stationary phase from ChIP-seq experiments (from Kahramanoglou et al 2010) | 3,2 |
|  | Putative H-NS target genes in stationary phase from ChIP-seq experiments (from Kahramanoglou et al 2010) | 2,4 |
|  | Genes overlapping with tsEPODs (transcriptionally silent extended protein occupancy domains) (from Vora et al 2009) | 2,2 |
| RH | Genes overlapping with heEPODs (highly expressed extended protein occupancy domains) (from Vora et al 2009) | 5,4 |
|  | Genes overlapping with tsEPODs (transcriptionally silent extended protein occupancy domains) (from Vora et al 2009) | 3,4 |

Table S6: **Only the subpopulation made of low-expression clones is consistent with the result of the whole population.** The data in this table refer to Z-scores for the first round high expression (RH) and low-expression (RL) FACS-sorted populations. The table reports the gene sets that result over-represented with Z-score $> +3$.

#### Supplementary Files

The Excel supplementary file SF1 *SF1_Suppl_ListColoniesNew.xlsx* contains a list of all the single insertion clonal strains analyzed in detail, including insertion coordinate and measured gene expression parameters (level and noise).

**Legend:**

Column A: Denotes the population where the clone comes from.

Column B: Describes the origin of the clone and which round of FACS the population came from.

Column C: The ID of the clonal colony (defined as its position in the 96-well plate, arranged column wise).

Column D: The coordinate position of the insertion in the genome, from sequencing. 'N' denotes the positions in the âĂIJrrnâĂİ regions (note that we do not know the location of exact insertion in very low expressing populations, since they are paired with different rrn operons such as 2728Kb , 4210 kb, 3426kb, 4169kb).

Column E: 'Y' denotes whether each clonal population underwent flow cytometry or not, to measure the CV, Mean and noise of the expression of GFP in each individual strain.

Column F: 'Y' denotes whether the fluorimeter measurements were made for this clone.

Column G: indicates the average protein concentration of individual colonies grown in Glucose at 37 degrees (n=3).

Column H: indicates the average protein concentration of individual colonies grown in Glucose + CAA at 37 degrees (n=3).

Column I: describes the average growth rate (doublings/hour) of individual colonies grown in Glucose at 37 degrees.

Column J: describes the average growth rate (Doublings/hour) of individual colonies grown in Glucose + CAA at 37 degrees.

Column K: describes the mean (Log) fluorescence measurements of individual colonies that underwent flow cytometry.

Column L: describes the CV (Log) fluorescence measurements of individual colonies that underwent flow cytometry.
